# Associations Between Neural Responses to Social Fairness and Financial Exploitation Vulnerability Vary With Age and Emotional Intelligence

**DOI:** 10.64898/2026.09.30.755119

**Authors:** Jacob M. Stanley, Daniel Sazhin, Chelsea Helion, Tania Giovannetti, Dominic Fareri, David V. Smith

## Abstract

Financial exploitation often arises in social contexts in which individuals must evaluate fairness, trust, and interpersonal information during financial decisions. We examined whether neural responses to fairness in social versus nonsocial contexts relate to financial exploitation vulnerability and whether these relationships vary with age, emotional intelligence, and affect. A total of 310 adults spanning ages 21–89 completed an fMRI Ultimatum Game in which they accepted or rejected offers from human and computer partners, along with measures of financial exploitation vulnerability, emotional intelligence, and affect. Participants were strongly sensitive to offer fairness, and individual differences in behavioral sensitivity to partner type were associated with social-relative anterior insula responses to unfairness. Greater baseline negative affect was associated with stronger anterior insula differentiation between social and nonsocial fairness. Financial exploitation vulnerability was associated with prefrontal responses to social-relative fairness in ways that depended on individual characteristics: age moderated its association with dorsomedial prefrontal cortex activation, whereas emotional intelligence moderated its association with dorsolateral prefrontal cortex activation. Whole-brain connectivity analyses further showed that age moderated associations between emotional intelligence and executive control network connectivity with posterior cingulate and right temporoparietal regions. These findings suggest that vulnerability to financial exploitation is reflected in how social context is integrated with affective and individual characteristics across adulthood.

## Introduction

Financial exploitation often begins as a social decision. A person must decide whether to trust a request, recognize that an exchange is unfair, or weigh a financial outcome against a relationship or persuasive appeal (Hanoch et al., 2025; Stanley & Smith, 2025). These judgments help explain why vulnerability cannot be reduced to chronological age. Across community samples, greater vulnerability has been associated with difficulties regulating emotion, greater persuadability, reduced sensitivity to trustworthiness, lower social support, and cognitive challenges (Lichtenberg et al., 2020; Yang et al., 2025). Neuroimaging studies of financially exploited older adults implicate socioemotional systems, including structural differences in anterior insula and resting-state connectivity involving insular, cingulate, and medial frontal regions (Spreng et al., 2017; Weissberger et al., 2020; Smith et al., 2026). Yet these studies have primarily characterized brain structure or intrinsic connectivity, leaving unclear whether vulnerability is expressed in neural responses during social-economic choice.

The Ultimatum Game (UG) provides a validated and well-characterized model for examining responses to disadvantageous social economic exchange. In the UG, a responder plays with a partner and accepts or rejects proposed divisions of money from an initial endowment. If the responder accepts, both the responder and partner receive their agreed upon proportion; however, if the responder rejects, both parties receive nothing. Behaviorally, studies of the UG have shown that people frequently reject highly unequal offers even when acceptance would increase their own payoff (Güth et al., 1982; Sanfey et al., 2003). Neuroimaging studies have shown that unfair offers recruit the anterior insula and anterior cingulate cortex, and decisions to reject them additionally engage regions involved in action selection and control (Gabay et al., 2014; Ră□ală & Sanfey, 2026). Fair treatment also carries positive value, recruiting reward-related circuitry even when monetary payoff is held constant (Tabibnia et al., 2008).

Comparing offers attributed to human and computer partners tests how these responses depend on the source of an offer (Sanfey et al., 2003). Social information can itself alter valuation: observing others’ choices changes the subjective utility of risky options, with this “other-conferred utility” represented in vmPFC and individual preference incongruence reflected in anterior cingulate and insular responses (Chung et al., 2015). Interactions among medial prefrontal, insular, and reward-related systems have been implicated in social connectedness and social evaluation (Kim and Sul, 2023). More broadly, neuroeconomics emphasizes that economic choice emerges from interacting valuation, affective, cognitive, and social processes, providing a framework for understanding how interpersonal context can alter the computations that guide decisions (Rangel et al., 2008; Ruff & Fehr, 2014; Dennison et al., 2022; Smith et al., 2026).

Responses to unfair exchange also vary with the characteristics and emotional state of the decision maker. Older and younger adults can differ in Ultimatum Game choices and in the balance of insular and prefrontal recruitment (Harlé and Sanfey, 2012; Bailey et al., 2013; Girardi et al., 2018). Experimentally induced sadness increases rejection of unfair offers, while cognitive reappraisal changes both choice and insular-prefrontal responses (Harlé and Sanfey, 2007; Grecucci et al., 2013). Emotional intelligence may be especially relevant because it reflects how people perceive, understand, and manage emotion in themselves and others, and higher ability emotional intelligence has been associated with lower susceptibility to investment fraud (Mueller et al., 2020). Age, affect, and emotional intelligence may therefore shape the neural meaning of an unfair offer and, through that route, help explain why some people are more vulnerable to financial exploitation.

We examined these relationships in 310 adults who completed an fMRI Ultimatum Game involving offers from human and computer partners, together with measures of financial exploitation vulnerability, trait emotional intelligence, and affect (Smith et al., 2024; Campbell and Lichtenberg, 2021; Cooper and Petrides, 2010; Watson et al., 1988). We asked three questions. First, how does fairness shape choices involving human and computer partners, and how is individual variation in sensitivity to partner type represented neurally? Second, how do age, emotional intelligence, and negative affect relate to neural sensitivity to social versus nonsocial fairness? Third, does this neural sensitivity relate to financial exploitation vulnerability, and are those associations conditioned by age or emotional intelligence?

## Methods

### Participants and Recruitment

We collected data from 310 adults (age range 21–89 years) recruited from the Temple University community and the surrounding Philadelphia area. This sample was drawn from a larger parent project designed to examine social decision making across the adult lifespan; the present study represents one of several investigations using data from this project. Sample size and exclusion criteria were pre-registered on OSF: https://osf.io/9ezhy/overview. Eligible participants were native English speakers with normal or corrected to normal vision and no history of neurological or psychiatric illness. All participants completed a health screening questionnaire and standard MRI safety screening prior to enrollment. Participants were told that one of the trials would be selected at random and added to their bonus payment. They were also informed that proposer partners would receive payment based on that random trial’s outcome, with any money won either going directly to the social partner or being returned to a pool of laboratory funds for outcomes with nonsocial partners. The study protocol was approved by the Temple University Institutional Review Board, and all participants provided written informed consent.

For the present analyses, participants were excluded as outliers for demonstrating poor neuroimaging data quality (as assessed with MRIQC). We removed runs using a boxplot threshold applied to the tsnr and fd_mean IQMs from MRIQC. Specifically, outlier runs were defined as runs with fd_mean values exceeding Q3 + 1.5 × IQR, as well as those with tsnr values lower than Q1 − 1.5 × IQR, where Q1 and Q3 are the 25th and 75th percentiles, respectively, and IQR = Q3 − Q1. We note that the tsnr for the multi-echo data were derived from echo-2 to increase compatibility with the single-echo data. In addition, runs where participants missed more than 25% of task trials were excluded. After all the exclusions, data from 258 participants were included in the analysis.

### Experimental Procedure

Data collection occurred over two visits. During Visit 1 (≈ 90 min), participants underwent a mock MRI scan to acclimate to scanner noise and the supine position. Visit 2 (≈ 120 min), scheduled within one week of Visit 1, began with practice trials of the Ultimatum Game (UG) on a laptop to minimize missing responses, followed by an fMRI run. After the scan, participants completed exploratory questionnaires.

### Individual-Difference Measures

Financial exploitation vulnerability was assessed once using the nine-item Financial Exploitation Vulnerability Scale–Short Form (FEVS-SF; Campbell & Lichtenberg, 2021). Each item was scored 0, 1, or 2 and summed (range = 0–18), with higher scores indicating greater vulnerability. Trait emotional intelligence was assessed once using the 30-item Trait Emotional Intelligence Questionnaire–Short Form (TEIQue-SF; Cooper & Petrides, 2010). Items were rated from 1 (completely disagree) to 7 (completely agree); negatively keyed items were reverse-scored, and the mean of all 30 items provided the global trait emotional-intelligence score (range = 1–7).

Affect was assessed at prescan and postscan time points surrounding the fMRI session using the 20-item Positive and Negative Affect Schedule (PANAS; Watson et al., 1988). Participants rated 10 positive- and 10 negative-affect adjectives from 1 to 5; each subscale was summed separately (range = 10–50), with higher scores indicating greater affect. The prescan negative-affect total indexed baseline negative affect, and the negative-affect change score was calculated as postscan minus prescan, so positive values indicated increased negative affect across the session. Analyses used dense-rank versions of the prescan and change scores, as described below.

### Tasks and Experimental Design

In the UG, participants are made offers by one of two different partners (a stranger or a computer) and decide whether to accept or reject the offer (Figure 1). When paired with a (social) stranger partner, participants were informed that their partner was a past participant that made real choices in past games and that their choices may similarly be used with future participants. Each stranger was experienced only once to maintain the one-shot design of the game. When paired with a (nonsocial) computer partner, participants were informed that their choices were decided by an algorithm, and the winnings would return to the general pool of lab funds. Participants are told that the partner received $16 or $32 and decided to share 5%, 10%, 25%, or 50% of this endowment with them. If they accept the offer, they both receive the money as the split proposes. However, if they reject the offer, neither the participant nor the partner will receive any money. Each trial consisted of the endowment phase (1 second), an interstimulus interval of (1 second), a decision phase of 3.25 s, and an intertrial interval of 1–7 s (M = 2.5 s).

**Figure 1.**
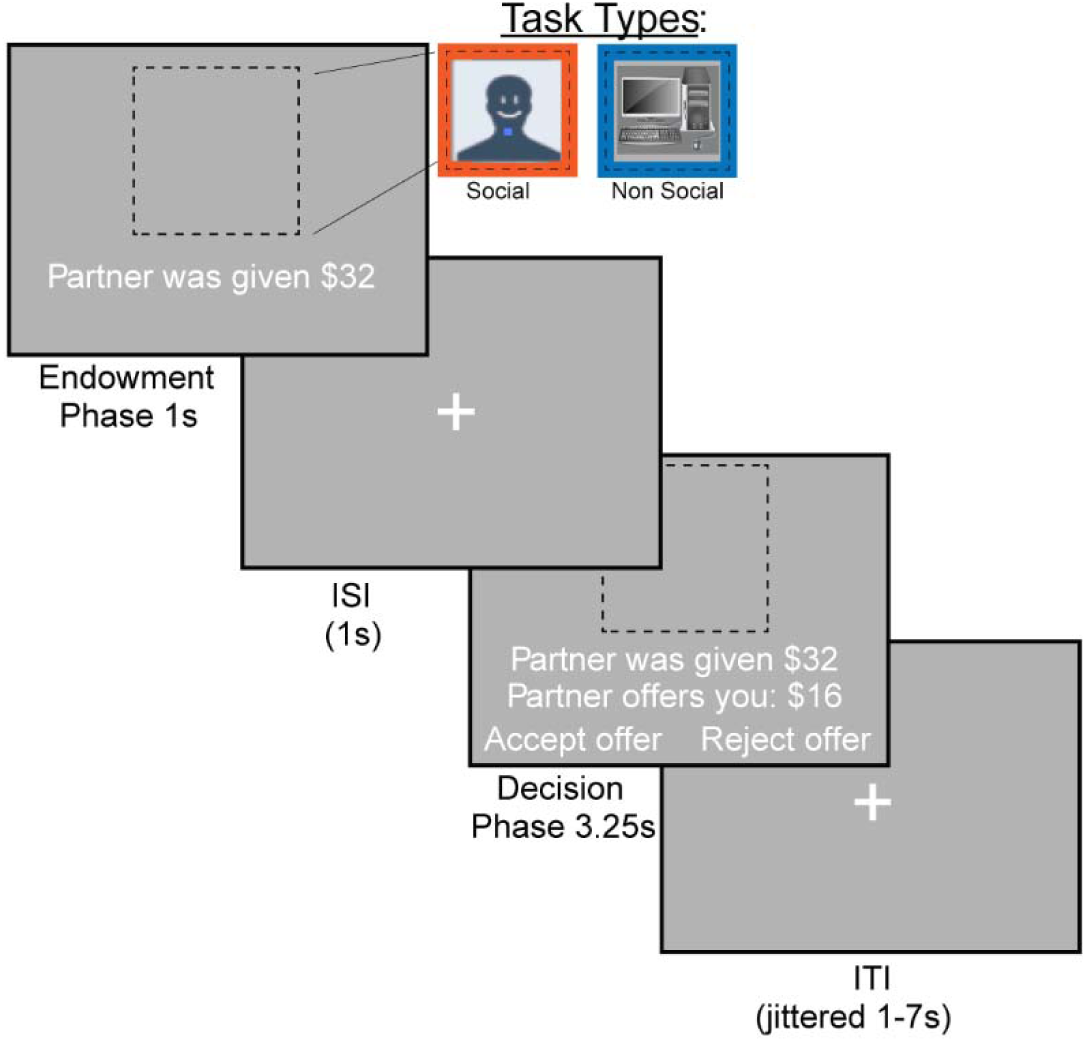
Task Timeline. Participants in the Ultimatum Game decided to accept or reject offers made by a social or nonsocial partner. If they rejected the offer, neither the participant nor their partner would make any money. Participants were offered between 5% and 50% of the initial endowment.

### Neuroimaging Preprocessing

Images were converted to Brain Imaging Data Structure format with HeuDiConv and preprocessed with fMRIPrep 24.1.1. Multi-echo functional runs underwent echo-time-dependent denoising with tedana, and MEDIC-derived B0 field maps were used for susceptibility-distortion correction. Preprocessed data were normalized to MNI152NLin6Asym standard space. Full acquisition, anatomical and functional preprocessing, denoising, distortion-correction, and nuisance-regressor details are provided in the Supplemental Methods.

### Neuroimaging Analysis

#### First-Level Activation Models

Analyses were conducted in FEAT (FSL v6.0.7; Jenkinson et al., 2012). Each run-level activation model focused on the offer phase and contained 11 task regressors: four offer-epoch constants for the crossed partner (social, nonsocial) and endowment ($32 high, $16 low) conditions; four corresponding parametric modulators for offer value; a response-time impulse at choice-feedback onset and its parametric modulator; and a missed-trial regressor. Valid offer epochs extended from partner-cue onset through choice feedback (5.25 s); missed epochs ended with the missed-decision phase. Offer values were demeaned within each partner-by-endowment condition and run, response times were demeaned across valid trials in each run, and FEAT orthogonalization was disabled. Task regressors were convolved with FSL’s double-gamma hemodynamic response function. The primary fairness contrast summed the four offer-value modulators; the social-relative fairness contrast assigned positive weights to social and negative weights to nonsocial modulators. Positive effects therefore reflected increasing responses with greater fairness, whereas negative effects reflected greater responses as offers became less fair. Models also estimated condition-specific constant and parametric effects, high-versus low-endowment effects, and separate social and nonsocial fairness effects. Nuisance regressors included six motion parameters, the first six aCompCor components, non-steady-state volumes, framewise displacement, and tedana components classified as non-BOLD. A 100-s high-pass filter, 5-mm full-width-at-half-maximum spatial smoothing, and FILM prewhitening were applied.

#### First-Level Connectivity Models

Generalized psychophysiological interaction (gPPI) models tested task-modulated connectivity for each seed region (McLaren et al., 2012). Each model retained the 11 task regressors, added the unconvolved seed time series, and included 11 interactions between that physiological series and the task regressors, yielding 23 explanatory variables before nuisance regressors. Because FEAT formed each interaction from an already convolved psychological regressor and the unconvolved physiological series, no additional convolution was applied. Connectivity contrasts followed the activation-model contrast structure.

For network PPI (nPPI) analyses, all 10 continuous network time courses were estimated jointly by spatial regression and demeaned. Each target-network model included the 11 task regressors, the unconvolved target network time course, the other nine network time courses as nuisance regressors, and 11 target-network-by-task interactions, yielding 32 explanatory variables before other nuisance regressors. Network interactions were not additionally convolved, and contrasts preserved the activation-model structure after insertion of the nuisance-network columns.

#### Within- and Group-Level Models

When both usable UGR runs were available, run-level estimates were combined within participant and session using fixed effects; a single usable run entered the group analysis directly. Of 258 participants, 219 contributed two-run estimates and 39 contributed one run (30 run 1; 9 run 2); sessions were not combined. Group analyses used FLAME 1 mixed-effects models (Woolrich et al., 2004). All designs included an intercept and mean-centered age, mean framewise displacement, temporal signal-to-noise ratio, scanner-software indicators, and sex indicators. Continuous predictors were centered, and interactions were products of their centered constituents. Six designs tested (1) FEVS, (2) FEVS × age, (3) FEVS × emotional intelligence, (4) emotional intelligence × age, (5) prescan negative affect or pre-to-postscan negative-affect change in separate models, and (6) social-minus-nonsocial behavioral fairness sensitivity. The two negative-affect predictors were dense-ranked. Each predictor and interaction was tested in both directions while retaining the constituent main effects and common covariates. Activation analyses used the overall and social-minus-nonsocial fairness contrasts; nPPI analyses used those contrasts and the target-network main-effect contrast (cope 18).

#### Regions, Networks, and Statistical Inference

Preregistered activation regions of interest (ROIs) were the anterior insula (AI), dorsal anterior cingulate cortex (dACC), and dorsolateral prefrontal cortex (dlPFC). The AI mask was the Neurosynth association-test map for “anterior insula” thresholded at Z = 10. The dACC mask was based on the Neurosynth association-test map for “dACC” thresholded at Z = 5.37, with its center represented by a 10-mm-radius sphere constrained to exclude supplementary motor cortex. The dlPFC mask comprised the middle frontal gyrus in the Harvard–Oxford atlas. The executive control network (ECN) was network map 7 from the 10-network intrinsic-connectivity decomposition reported by Smith et al. (2009). Whole-brain maps were evaluated with a cluster-forming threshold of Z > 3.1 and Gaussian-random-field cluster-extent correction at p < .05 (Worsley, 2001), thereby controlling familywise error across the whole-brain search volume. Thresholded maps were visualized on MNI152NLin6Asym standard-space anatomy, and peak locations were reported in MNI coordinates.

#### Preregistration and Analytic Deviations

Hypotheses, the sample-size target, exclusion criteria, and planned analyses were preregistered on OSF (https://osf.io/9ezhy/overview). The final analyses deviated from the preregistered plan in several respects. First, behavioral sensitivity to fairness was quantified using participant-specific slopes across offer levels rather than acceptance rates restricted to unfair offers. Second, prescan negative affect and pre-to-postscan change in negative affect were converted to dense ranks to reduce leverage from their strongly right-skewed, tied distributions while preserving score order; this transformation was not preregistered. Third, although the preregistration placed the H2 and H3 individual-difference predictors in a common model, the final analyses evaluated the age × emotional-intelligence hypotheses and the baseline- and change-in-negative-affect hypotheses in separate models. This separation isolated each hypothesis and avoided combining correlated predictors and interaction terms in an overparameterized model. Finally, the preregistration was internally inconsistent for H4.3: the hypothesis identified dlPFC activation and ECN–dACC connectivity as the neural predictors of interest, whereas the corresponding analysis description named different neural measures. We followed the measures stated in H4.3 and tested whether dlPFC activation and ECN–dACC connectivity explained additional variance in financial exploitation vulnerability beyond behavioral and individual-difference predictors. As preregistered, a priori ROI analyses were followed by whole-brain analyses of the same contrasts.

## Results

To address our three primary questions, we first examined behavior in the Ultimatum Game (UG). As illustrated in Figure 1, participants accepted or rejected monetary offers made by either a social or nonsocial partner, allowing us to quantify sensitivity to offer fairness and determine whether this sensitivity varied as a function of partner type. We then tested whether individual differences in age, emotional intelligence, negative affect, and financial exploitation vulnerability were associated with neural responses to fairness in social versus nonsocial contexts. These analyses included preregistered region-of-interest tests followed by secondary whole-brain analyses to characterize effects beyond the hypothesized regions. We next examined whether individual differences in behavioral sensitivity to partner type were reflected in fairness-related neural responses. Finally, we tested whether age, emotional intelligence, affect, and financial exploitation vulnerability were related to task-modulated functional connectivity during social-relative fairness processing.

### Acceptance Rates Increase with Offer Fairness

Consistent with prior work, the UG tasks participants to accept or reject monetary offers from a partner’s endowment, which provides a lens through which to examine fairness processing by comparing acceptance rates across levels of offer equity (Figure 1).

Behavioral responses in the Ultimatum Game were analyzed to quantify acceptance rates as a function of offer fairness and partner type. Trial-level accept/reject decisions were converted to a binary variable (Accept = 1, Reject = 0). For each participant, we computed mean acceptance rates separately for each fairness level (5%, 10%, 25%, 50% of the endowment) and each partner condition (social vs. nonsocial). These subject-level acceptance rates served as the dependent measure in a two-way repeated-measures ANOVA with fairness (4 levels) and partner (2 levels) as within-subject factors.

We hypothesized that overall participants would reject unfair offers more often than fair offers. The preregistered analysis revealed a robust main effect of fairness, *F*(3, 771) = 355.58, *p* < .001, η² = .58, indicating that acceptance rates increased strongly as offers became more equitable. In contrast, there was no main effect of partner, *F*(1, 257) = 0.03, *p* = .863, and no interaction of fairness and partner, *F*(3, 771) = 0.13, *p* = .94. Thus, participants responded robustly to fairness of the offer but did not differentiate between social and nonsocial proposers at the behavioral level (Figure 2).

**Figure 2.**
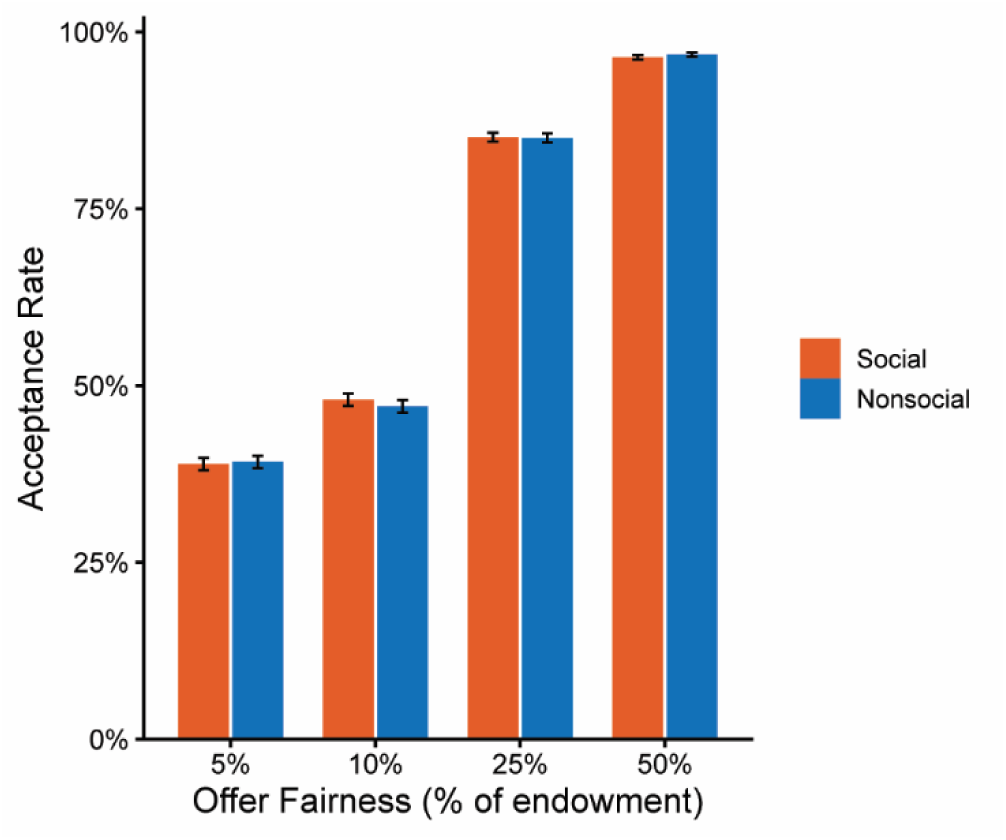
Acceptance Rates as a Function of Offer Fairness and Partner Type. Mean acceptance rates in the Ultimatum Game across four levels of offer fairness for social and nonsocial partners. Error bars represent ±1 standard error of the mean.

We hypothesized that older adults would accept unfair offers more often than younger adults. To test this hypothesis, we quantified each participant’s behavioral sensitivity to fairness from trial-level choices. Acceptance was modeled with a binomial generalized linear mixed-effects model that included offer fairness as a continuous fixed effect and allowed both the intercept and the fairness slope to vary across participants. We fit separate models to all trials, social-partner trials, and nonsocial-partner trials. For each model, a participant’s fairness slope was calculated by adding the population-level fairness coefficient to that participant’s random-slope deviation. We then calculated social-relative fairness sensitivity by subtracting the nonsocial slope from the social slope; larger values indicated a steeper increase in acceptance with fairness for human than computer partners. Additional derivation details are provided in the Supplemental Methods. We fit a preregistered linear model testing the effect of age on slope scores combined across partner type, *F*(1, 256) = 0.66, *p* = .42. There was no relationship found between age and behavioral sensitivity to offer fairness.

We hypothesized that emotional intelligence would moderate the relation between age and fairness, with lower EI scores rejecting offers more often in the social condition. To test this hypothesis, we fit a preregistered linear model testing the effect of age, emotional intelligence, and their interaction on difference slope scores, *F*(3, 254) = 0.24, *p* = .87. We found no interaction between age and emotional intelligence on social minus nonsocial difference slope scores, nor did we find any main effects of age or EI.

### Age and Emotional Intelligence Relate to Financial Exploitation Vulnerability

We hypothesized that participants that rejected unfair offers less often in social versus nonsocial contexts would be older in age, have lower EI scores, and will show greater vulnerability for financial exploitation (FEVS). To test this, we conducted a preregistered analysis which regressed financial exploitation vulnerability (FEVS) scores on participants’ social - nonsocial difference in behavioral sensitivity to fairness, age, and emotional intelligence. The overall model was significant, F(3, 254) = 15.88, p < .001, *R*² = .158. Lower EI significantly predicted greater financial exploitation vulnerability, *b* = −1.71, *t* = −4.82, *p* < .001, consistent with our hypothesis. However, age was significantly negatively associated with FEVS scores, *b* = −0.058, *t* = −3.96, *p* < .001, indicating that older participants reported lower financial exploitation vulnerability. These relationships are illustrated in Figure 3. The social - nonsocial difference in behavioral sensitivity to fairness was not significantly associated with FEVS scores, *b* = 0.049, *p* = .12.

**Figure 3.**
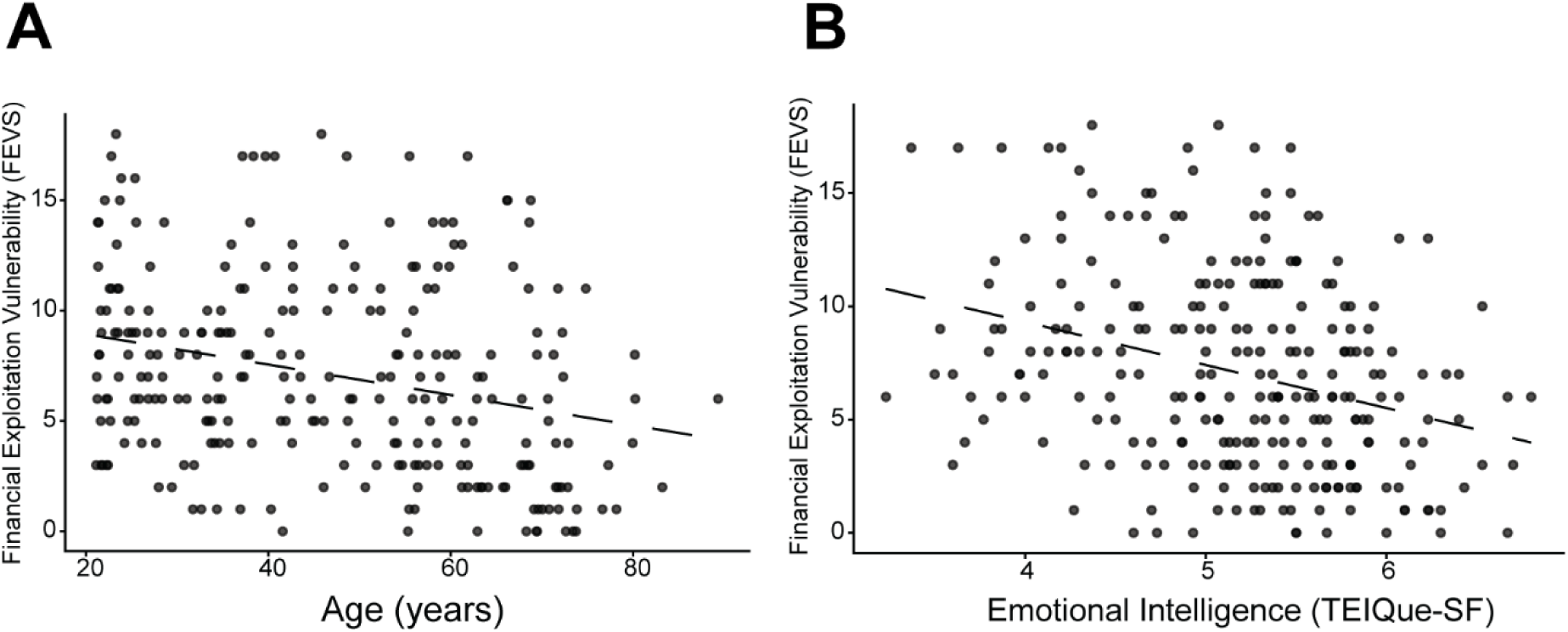
Financial Exploitation Vulnerability as a Function of Age and Emotional Intelligence. Scatterplots show associations between Financial Exploitation Vulnerability Scale (FEVS) scores and **(A)** age and **(B)** emotional intelligence (TEIQue-SF). Each point represents an individual participant, and dashed lines represent fitted linear regression lines. Greater age and higher emotional intelligence were each associated with lower financial exploitation vulnerability.

### Negative Affect Is Associated with Anterior Insula Sensitivity to Social Fairness

We hypothesized that increased dorsal Anterior Cingulate Cortex (dACC), Anterior Insula (AI), and dorsolateral prefrontal cortex (dlPFC) responses to unfair versus fair offers in the social versus nonsocial condition would be modulated by EI and that this effect will be attenuated with increasing age. To test this, we conducted preregistered analyses examining whether emotional intelligence (EI) and age interacted to predict BOLD responses within the dACC, AI, and dlPFC ROIs for the social versus nonsocial fairness contrast.

Our preregistered hypotheses were not supported. The overall model predicting dACC responses was not significant, *F*(3, 254) = 0.76, *p* = .516. The predicted EI × age interaction was also not significant, *b* = −0.0051, *p* = .234. Neither EI, *b* = 0.263, *p* = .213, nor age, *b* = 0.024, *p* = .292, was significantly associated with dACC responses. For the AI, the overall regression model was not significant, *F*(3, 254) = 1.22, *p* = .304. The EI × age interaction was also not significant, *b* = −0.0038, *p* = .347. Neither EI, *b* = 0.057, *p* = .774, nor age, *b* = 0.020, *p* = .332, significantly predicted AI responses. For the dlPFC, the overall regression model was not significant, *F*(3, 254) = 0.83, *p* = .478. The EI × age interaction was not significant, *b* = −0.0005, *p* = .862. Neither EI, *b* = −0.038, *p* = .799, nor age, *b* = 0.005, *p* = .734, was significantly associated with dlPFC responses. Thus, there was no evidence that the relationship between EI and either dACC, AI, or dlPFC responses was attenuated with increasing age in these preregistered ROI analyses. A follow-up, whole brain, cluster-corrected analysis was conducted and found no significant clusters where there was a significant interaction between EI and age.

We hypothesized that AI responses to unfair versus fair offers in the social versus nonsocial condition will be positively associated with baseline negative affect and with increases in negative affect. To test this, we examined baseline negative affect and change in negative affect as separate predictors of anterior insula (AI) responses in the social versus nonsocial constrast. Higher baseline negative affect was significantly associated with greater AI responses, *b* = 0.041, *t*(256) = 2.89, p = .004. The overall model was significant, *F*(1, 256) = 8.35, *p* = .004, *R*² = .032. This relationship is illustrated in Figure 4. In a separate model, change in negative affect from pre- to post-scan was not significantly associated with AI responses, *b* = −0.025, *p* = .190. This model was not significant, *F*(1, 256) = 1.73, *p* = .190. Thus, as shown in Figure 4, greater baseline negative affect was associated with greater AI responses as predicted; however, increases in negative affect were not significantly associated with AI responses.

**Figure 4.**
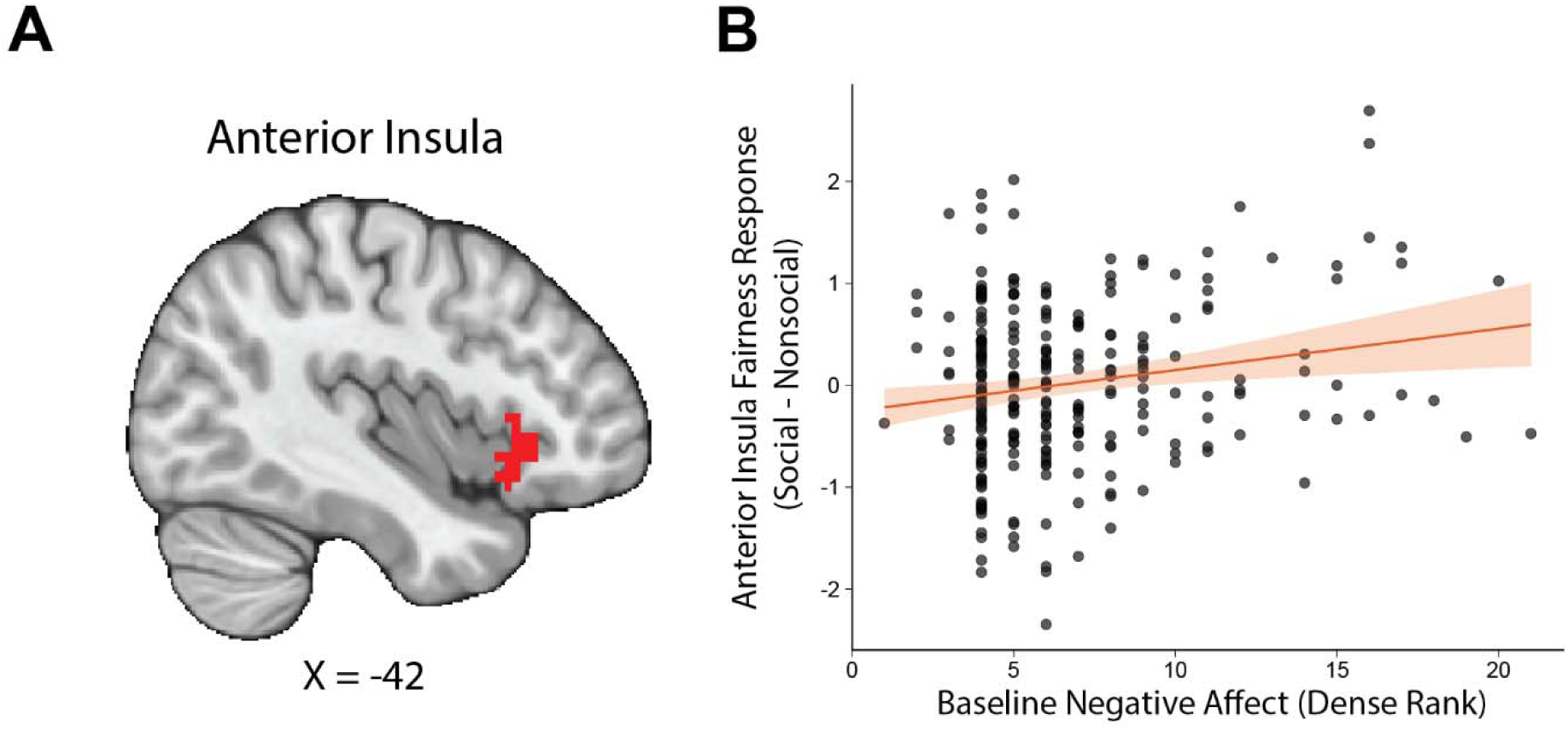
Baseline Negative Affect Is Associated With Anterior Insula Responses to Fairness. **(A)** Anterior insula (AI) region of interest used to quantify fairness-related activation. **(B)** Association between baseline negative affect and AI responses to fairness in social versus nonsocial contexts. Each point represents an individual participant. The solid line represents the fitted linear regression and the shaded region represents the 95% confidence interval.

### Financial Exploitation Vulnerability Relates to Prefrontal Fairness Responses Across Individual Differences

Consistent with our preregistration, we conducted secondary whole-brain analyses examining whether individual differences in financial exploitation vulnerability were associated with activation tracking fairness in social versus nonsocial contexts. These analyses identified a significant FEVS × age interaction in the dorsomedial prefrontal cortex (dmPFC; peak MNI = **-**1.9, 44.1, 32, peak Z = 4.32, *k* = 47, cluster-corrected *p* < .001). To characterize this interaction, mean activation was extracted from the significant cluster. The extracted-signal model was significant, *F*(3, 254) = 7.40, *p* < .001, *R*² = .080, and showed a positive FEVS × age interaction, *b* = 0.00349, *t*(254) = 4.54, *p* < .001. Johnson–Neyman analysis indicated that greater FEVS scores were associated with lower dmPFC activation tracking social fairness among participants younger than 45.1 years, were not significantly associated with dmPFC activation between ages 45.1 and 62.7 years, and were associated with greater dmPFC activation among participants older than 62.7 years. Thus, the relationship between financial exploitation vulnerability and dmPFC responses reversed across age (Figure 5).

**Figure 5.**
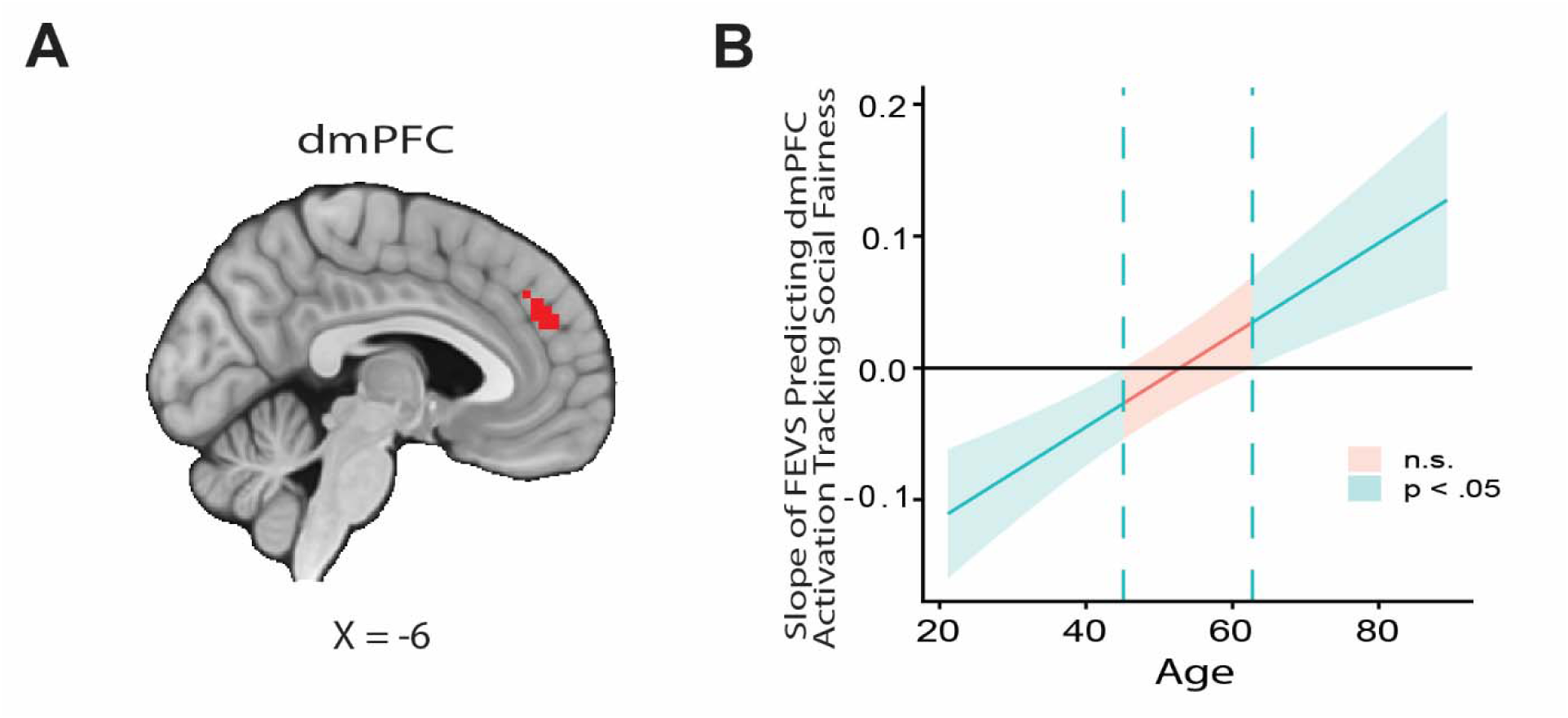
Age Moderates the Association Between Financial Exploitation Vulnerability and dmPFC Responses to Social Fairness. **(A)** Whole-brain analysis identified a dorsomedial prefrontal cortex (dmPFC) cluster showing a significant interaction between Financial Exploitation Vulnerability Scale (FEVS) scores and age for activation tracking fairness in social versus nonsocial contexts. **(B)** Johnson–Neyman plot showing the estimated slope of FEVS predicting dmPFC activation across age. Shaded regions indicate 95% confidence intervals; dashed vertical lines indicate Johnson–Neyman significance boundaries.

A second whole-brain analysis identified a significant FEVS × emotional intelligence interaction in the dorsolateral prefrontal cortex (dlPFC; peak MNI = 38.6, 33.3, 34.9, peak Z = 4.84, *k* = 42, cluster-corrected *p* = .001). Characterization of the extracted cluster signal showed a significant overall model, *F*(3, 254) = 9.52, *p* < .001, *R*² = .101, with a negative FEVS × EI interaction, *b* = −0.0993, *t*(254) = −5.08, *p* < .001. Johnson–Neyman analysis indicated that greater FEVS scores were associated with greater dlPFC activation tracking social fairness at EI scores below 4.53, were not significantly associated with dlPFC activation between EI scores of 4.53 and 5.16, and were associated with lower dlPFC activation at EI scores above 5.16. Thus, the relationship between financial exploitation vulnerability and dlPFC responses varied substantially as a function of emotional intelligence (Figure 6).

**Figure 6.**
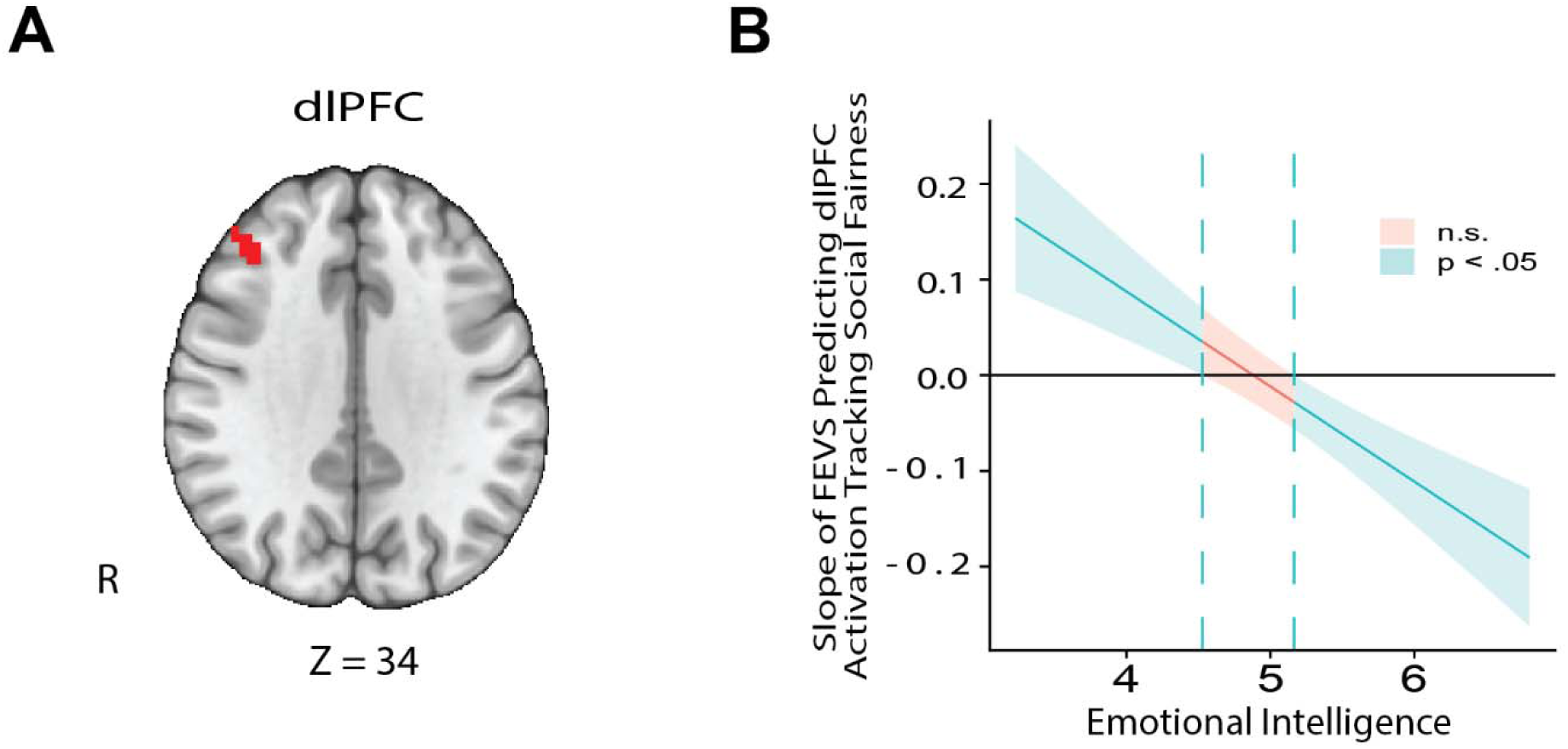
Emotional Intelligence Moderates the Association Between Financial Exploitation Vulnerability and dlPFC Responses to Social Fairness. **(A)** Whole-brain analysis identified a dorsolateral prefrontal cortex (dlPFC) cluster showing a significant interaction between FEVS scores and emotional intelligence for activation tracking fairness in social versus nonsocial contexts. **(B)** Johnson–Neyman plot showing the estimated slope of FEVS predicting dlPFC activation across levels of emotional intelligence. Shaded regions indicate 95% confidence intervals; dashed vertical lines indicate Johnson–Neyman significance boundaries.

### Behavioral Sensitivity to Social Fairness Tracks Anterior Insula Responses

We next conducted an exploratory whole-brain analysis to determine whether individual differences in behavioral sensitivity to partner type were reflected in neural responses to the same manipulation. Behavioral partner-type sensitivity was quantified as the social-minus-nonsocial difference in participants’ fairness slopes, such that larger values reflected greater behavioral sensitivity to fairness when interacting with human relative to computer partners. These difference slopes were entered as a covariate in a whole-brain model of the social-minus-nonsocial contrast in fairness-related activation.

Greater social-relative behavioral sensitivity to fairness was associated with a more negative social-minus-nonsocial fairness-modulation effect in the anterior insula; (peak MNI = 30.5, 30.6, -3.69; peak Z = 3.98; k = 30; cluster-corrected p = .011). Because the parametric modulator increased with offer fairness, a more negative modulation effect reflects relatively greater anterior insula engagement as offers became less fair. Thus, participants who behaviorally differentiated more strongly between social and nonsocial partners as a function of fairness also showed stronger social-relative anterior insula modulation by unfairness (Figure 7).

**Figure 7.**
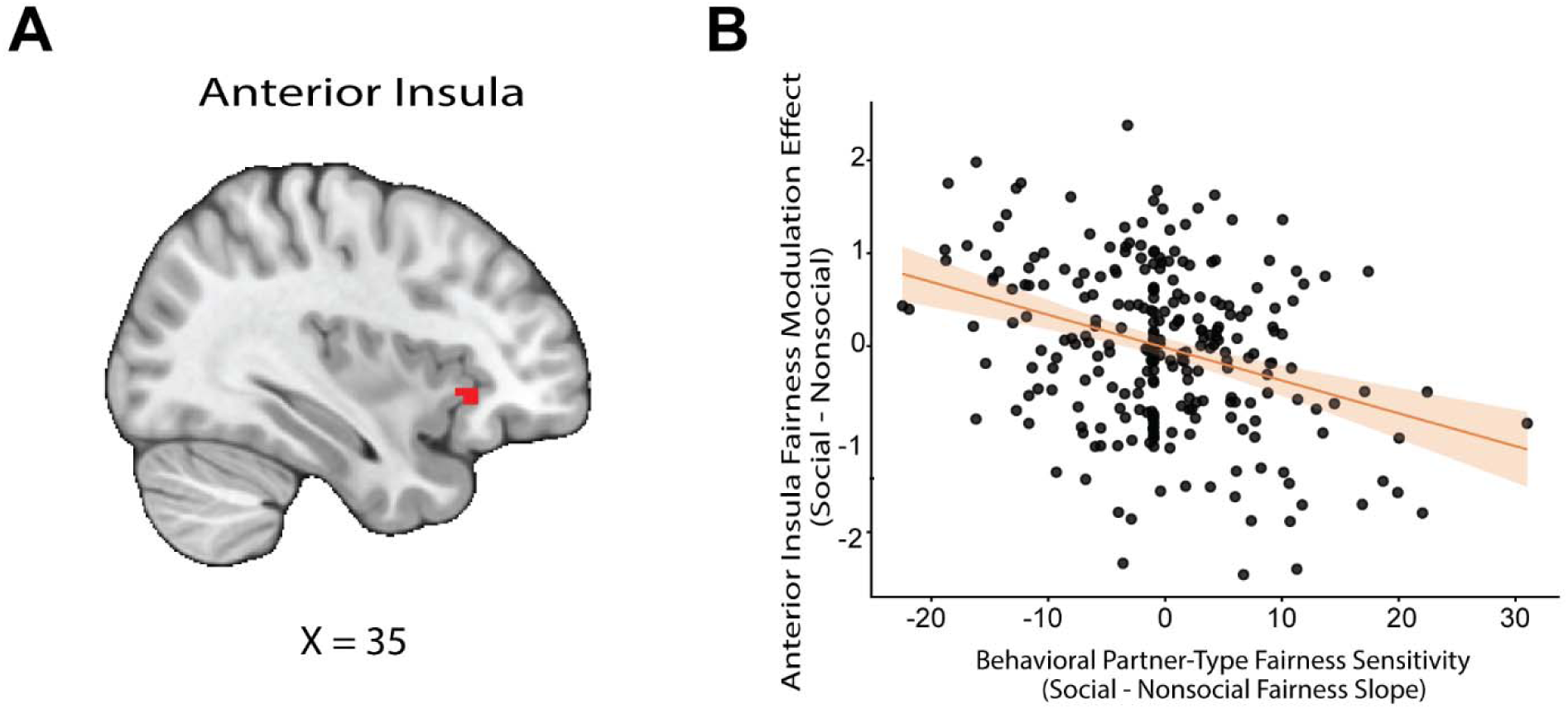
Association Between Behavioral Partner-Type Sensitivity and Anterior Insula Fairness Modulation. **(A)** Whole-brain analysis identified a cluster in the right anterior insula in which the social-minus-nonsocial fairness-modulation effect was associated with individual differences in behavioral partner-type sensitivity. **(B)** Scatterplot illustrating the association between behavioral partner-type fairness sensitivity, quantified as the social-minus-nonsocial difference in fairness slopes, and the extracted anterior insula fairness-modulation effect. Larger behavioral difference scores indicate greater sensitivity to fairness for social relative to nonsocial partners. The solid line represents the fitted linear regression and the shaded region represents the 95% confidence interval.

### Age Shapes Emotional Intelligence–Related Connectivity During Social Fairness

We hypothesized that ECN–dACC connectivity during unfair versus fair offers in social versus nonsocial contexts would be moderated by emotional intelligence (EI), and that this effect would be enhanced with increasing age. To test this hypothesis, we examined whether EI and age interacted to predict ECN–dACC connectivity for the social versus nonsocial fairness contrast. The overall regression model was not significant, *F*(3, 254) = 1.03, *p* = .378. Critically, the predicted EI × age interaction was not significant, *b* = −0.00049, *p* = .877. Neither EI, *b* = −0.015, *p* = .925, nor age, *b* = 0.006, *p* = .697, was significantly associated with ECN–dACC connectivity. Thus, there was no evidence that the relationship between EI and ECN–dACC connectivity varied as a function of age.

We hypothesized that AI–ECN connectivity during unfair versus fair offers in social versus nonsocial contexts would be associated with baseline negative affect and with increases in negative affect. To test this hypothesis, baseline negative affect and change in negative affect were examined as separate predictors of AI–ECN connectivity. Baseline negative affect was not significantly associated with AI–ECN connectivity, *b* = 0.0024, *p* = .802. The overall model was not significant, *F*(1, 256) = 0.06, *p* = .802. In a separate model, change in negative affect was also not significantly associated with AI–ECN connectivity, *b* = −0.0067, *p* = .609. This model was likewise not significant, *F*(1, 256) = 0.26, *p* = .609. Thus, neither baseline negative affect nor changes in negative affect were associated with AI–ECN connectivity.

We hypothesized that older adults showing the least differentiation in the dlPFC between unfair and fair offers in social versus nonsocial contexts will demonstrate the highest risk for financial exploitation. To test this, we examined whether age moderated the association between dlPFC responses to the social versus nonsocial fairness contrast and financial exploitation vulnerability (FEVS). The overall regression model was significant, *F*(3, 254) = 6.92, *p* < .001, *R*² = .076. However, the predicted dlPFC × age interaction was not significant, *b* = −0.014, *p* = .819, indicating no evidence that the association between dlPFC differentiation and financial exploitation vulnerability varied with age. The main effect of dlPFC response was also not significant, *b* = 0.379, *p* = .900. As reported earlier, age was significantly negatively associated with FEVS scores.

We hypothesized that older adults showing less differential ECN connectivity with the dACC when experiencing unfair versus fair offers in a social versus nonsocial context will demonstrate the highest risk for financial exploitation. To test this, we examined whether age moderated the association between ECN–dACC connectivity for the social versus nonsocial fairness contrast and financial exploitation vulnerability (FEVS). The overall regression model was significant, *F*(3, 254) = 8.06, *p* < .001, *R*² = .087. However, the predicted ECN–dACC connectivity × age interaction was not significant, *b* = −0.040, *p* = .101, indicating no evidence that the association between ECN–dACC connectivity and financial exploitation vulnerability differed as a function of age.

We hypothesized that dlPFC activation and ECN-dACC connectivity will be better predictors of risk for financial exploitation than age and social functioning. To test this, we used hierarchical regression to examine whether dlPFC activation and ECN–dACC connectivity explained additional variance in financial exploitation vulnerability beyond behavioral and individual-difference. The baseline model including social–nonsocial behavioral sensitivity to fairness, age, and emotional intelligence (EI) was significant, *F*(3, 254) = 15.88, *p* < .001, *R*² = .158. In this model, older age, *b* = −0.058, *t* = −3.96, *p* < .001, was associated with lower financial exploitation vulnerability, lower EI, *b* = −1.71, *t* = −4.82, *p* < .001, was associated with greater financial exploitation vulnerability, and behavioral sensitivity to fairness was not significant, *b* = 0.049, *p* = .115.

Adding dlPFC activation and ECN–dACC connectivity produced a significant overall model, *F*(5, 252) = 10.22, *p* < .001, *R*² = .169. However, the addition of the two brain measures did not significantly improve model fit relative to the baseline model, Δ*R*² = .011, *F*(2, 252) = 1.62, *p* = .201. Neither dlPFC activation, *b* = −0.72, *p* = .092, nor ECN–dACC connectivity, *b* = 0.24, *p* = .556, was a significant unique predictor of FEVS scores.

Following the a priori ROI connectivity analyses, whole-brain analyses examined whether age moderated the association between emotional intelligence and ECN connectivity tracking fairness in social versus nonsocial contexts. A significant EI × age interaction was observed for connectivity between the ECN and posterior cingulate cortex (PCC; peak MNI = -15.4, -50.4, 32, peak Z = 4.38, *k* = 15, cluster-corrected *p* = .035). Characterization of the extracted cluster signal showed a significant overall model, *F*(3, 254) = 10.70, *p* < .001, *R*² = .112, and a positive EI × age interaction, *b* = 0.0185, *t*(254) = 4.37, *p* < .001. Johnson–Neyman analysis indicated that higher EI was associated with lower PCC–ECN connectivity tracking social fairness among participants younger than 51.2 years. This association was not significant between ages 51.2 and 74.7 years, whereas among participants older than 74.7 years, higher EI was associated with greater PCC–ECN connectivity. Thus, the relationship between emotional intelligence and PCC–ECN connectivity reversed across age (Figure 8).

**Figure 8.**
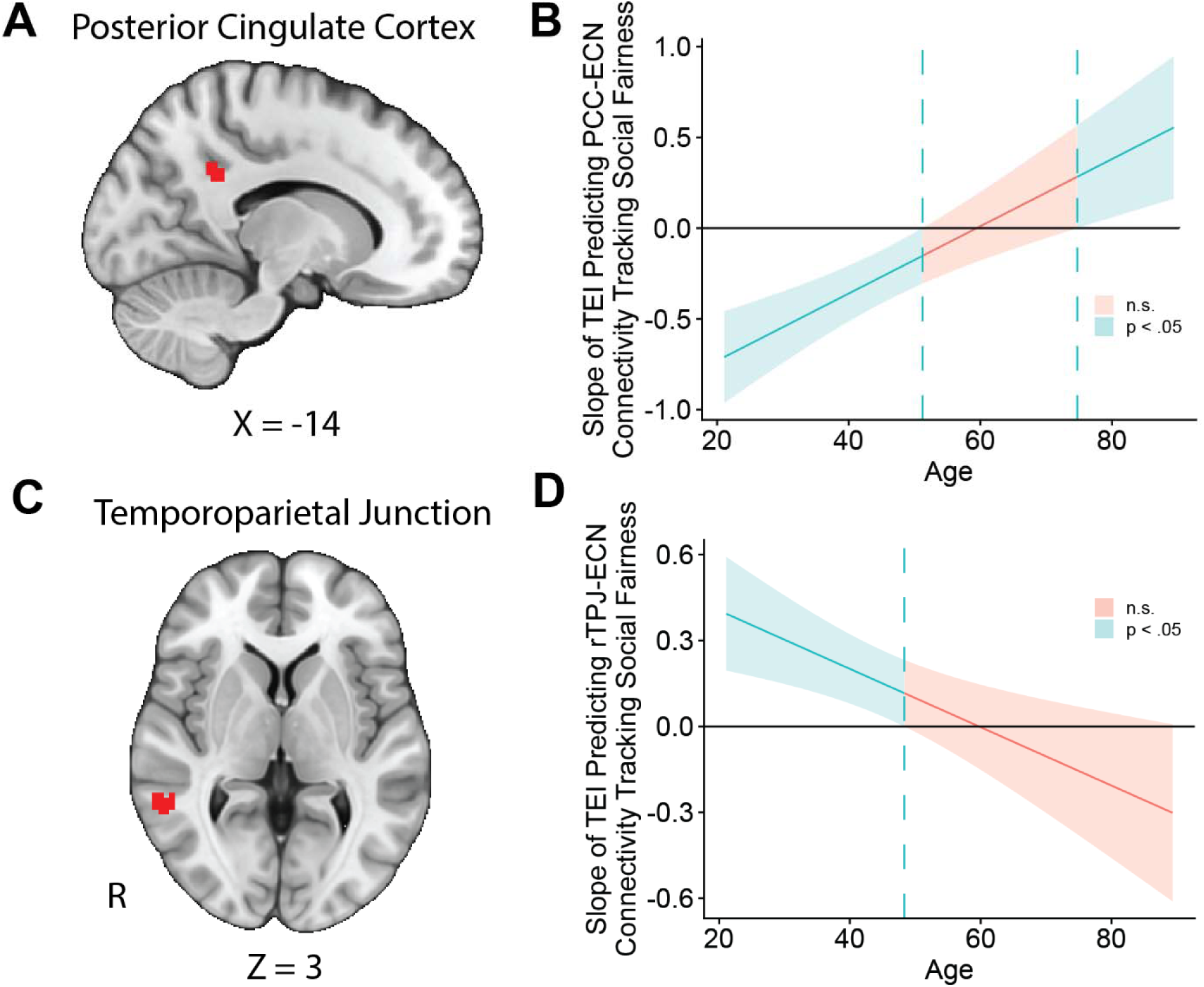
Age Moderates the Association Between Emotional Intelligence and ECN Connectivity During Social Fairness. **(A)** Whole-brain analysis identified a posterior cingulate cortex (PCC) cluster showing a significant emotional intelligence (EI) × age interaction in executive control network (ECN) connectivity tracking fairness in social versus nonsocial contexts. **(B)** Johnson–Neyman plot showing the estimated association between EI and PCC–ECN connectivity across age. **(C)** Whole-brain analysis identified a right temporoparietal junction (rTPJ) cluster showing a significant EI × age interaction in ECN connectivity. **(D)** Johnson–Neyman plot showing the estimated association between EI and rTPJ–ECN connectivity across age. Shaded regions indicate 95% confidence intervals; dashed vertical lines indicate Johnson–Neyman significance boundaries.

A second significant EI × age interaction was observed for connectivity between the ECN and right temporoparietal junction (rTPJ; peak MNI = 54.8, -47.7, 5.22, peak Z = 4.88, *k* = 25, cluster-corrected *p* = .002). The extracted-signal model was significant, *F*(3, 254) = 5.72, *p* < .001, *R*² = .063, with a negative EI × age interaction, *b* = −0.0102, *t*(254) = −3.05, *p* = .003. Johnson–Neyman analysis indicated that higher EI was associated with greater rTPJ–ECN connectivity tracking social fairness among participants younger than 48.3 years. Above age 48.3, the association was no longer statistically significant within the observed age range. Although the estimated association became increasingly negative with age, the second Johnson–Neyman boundary occurred at age 91.1, beyond the oldest participant in the sample; therefore, there was no observed age range in which the negative association was statistically significant (Figure8).

## Discussion

We used a social and nonsocial version of the Ultimatum Game to examine whether the neural processing of fairness helps explain financial exploitation vulnerability across adulthood. Fairness strongly organized participants’ choices, while average acceptance rates were similar for human and computer partners. Beneath that group-level similarity, individual differences in behavioral sensitivity to partner type corresponded to anterior insula responses to unfairness. Baseline negative affect also tracked the insular distinction between social and nonsocial fairness. At the network level, age altered how emotional intelligence related to executive control network connectivity with posterior cingulate and temporoparietal regions. Financial exploitation vulnerability was associated with dorsomedial and dorsolateral prefrontal responses in ways that depended on age and emotional intelligence. Together, these findings place vulnerability within the interaction of social context, affect, and individual characteristics during financial choice.

Our first question concerned how fairness guides choices with human and computer partners and how such influences are represented neurally. Participants were far more likely to accept offers as they became equitable, confirming that the task produced a strong and graded behavioral response to fairness. At the group level, this response was comparable across partner types. However, our analysis of individual variation relating fairness related behavior and neural response revealed that participants whose acceptance behavior was more sensitive to fairness with human than computer partners also showed greater social-relative anterior insula engagement as offers became unfair. The behavioral and neural measures therefore supported the anterior insula as expressing individual differences in social-relative fairness sensitivity which reflected acceptance behavior differences between partner types, even though the average behavioral curves overlapped.

This anterior insula association extends a substantial literature linking the region to disadvantageous inequity and the rejection of unfair offers (Sanfey et al., 2003; Nitsch et al., 2022). In the present study, the insular response was strongest among people whose choices changed most when the same fairness information was embedded in a social exchange. Exploitative solicitations are typically embedded in persuasive social interactions that can leverage trust, urgency, obligation, and other psychological vulnerabilities (Hanoch et al., 2025; Stanley & Smith, 2025). Recent neuroeconomic work likewise situates financial exploitation among maladaptive decision-making processes in older adults (Wyngaarden et al., 2026). Individual sensitivity to the social source of an offer may thus be an important bridge between laboratory measures of fairness and the interpersonal settings in which financial vulnerability becomes consequential.

Our second question asked how age and emotional functioning shape neural sensitivity to social fairness. Participants who entered the task with higher negative affect showed a greater anterior insula distinction between social and nonsocial fairness. This finding complements evidence that sadness and reappraisal alter Ultimatum Game behavior and insular-prefrontal responses (Harlé and Sanfey, 2007; Grecucci et al., 2013). It also connects an ongoing affective state to the way social context is represented during economic choice: negative affect may heighten the motivational significance of an unfair offer when it comes from another person.

Age and emotional intelligence were most clearly expressed in interactions between large-scale networks. Among younger adults, higher emotional intelligence was associated with lower coupling between the executive control network and posterior cingulate cortex during social-relative fairness processing; among the oldest participants, this association reversed. Higher emotional intelligence was also associated with greater executive control network coupling with right temporoparietal junction in younger adults, and this relationship weakened across age. Posterior cingulate cortex supports internally directed cognition, whereas right temporoparietal junction contributes to representing other people’s perspectives (Spreng et al., 2010; Schurz et al., 2014). Their age-dependent coupling with the executive control network suggests that emotional intelligence relies on a changing balance of internally directed cognition, social inference, and control across adulthood.

The third question addressed the link between neural sensitivity to social fairness and financial exploitation vulnerability. Vulnerability was associated with dorsomedial prefrontal fairness responses differently across age: greater vulnerability accompanied lower social-relative responses among younger adults and higher responses among older adults. Emotional intelligence similarly changed the relationship between vulnerability and dorsolateral prefrontal activation. At lower emotional intelligence, greater vulnerability accompanied higher social-relative dorsolateral prefrontal responses; at higher emotional intelligence, it accompanied lower responses. These crossovers show that the relevance of prefrontal recruitment on financial exploitation vulnerability depends on the person in whom it is observed. Engagement of the same prefrontal brain regions may be either beneficial or maladaptive depending on the age and emotional intelligence of the decision maker.

The prefrontal findings extend previous structural and resting-state observations into active decision making (Spreng et al., 2017; Weissberger et al., 2020). Medial prefrontal systems contribute to social inference and the construction of subjective value, while dorsolateral prefrontal cortex supports the implementation of fairness-related choices (Hampton et al., 2006; Jenkins et al., 2008; Bartra et al., 2013; Knoch et al., 2006). Our results show that vulnerability is reflected in these systems during a choice that combines material outcomes with information about another agent. The changing direction of the associations across age and emotional intelligence further suggests that vulnerability can arise through different configurations of social contexts and individual differences. This provides a neural complement to evidence that financial exploitation risk is heterogeneous and shaped by psychological and relational characteristics (Lichtenberg et al., 2020; Yang et al., 2025).

## Acknowledgments

We thank Cooper Sharp and Melanie Kos for assistance with data collection, preprocessing, and analysis, and Avi Dachs, Derrick Dwamena, Shenghan Wang, and Jamie-Nicole Luistro for assistance with data collection. This work was supported by National Institutes of Health grant R01-AG067011. Data are publicly available on OpenNeuro (https://openneuro.org/datasets/ds005123/versions/1.1.3), and analysis code and selected derivative outputs are available on GitHub (https://github.com/DVS-Lab/rf1-ugr-fevs).

