## Supplemental Methods for "Associations Between Neural Responses to Social Fairness and Financial Exploitation Vulnerability Vary With Age and Emotional Intelligence"

The imaging and behavioral data used in the present analyses were collected as part of a larger project described in Smith et al. (2024) and are publicly available on OpenNeuro (https://openneuro.org/datasets/ds005123/versions/1.1.3).

### Behavioral Analyses

Trial-level choices were coded as Accept = 1 and Reject = 0. Missed trials and trials from excluded runs did not enter the models. Because offers had to use whole-dollar amounts, the displayed proportions in the nominal 5% and 10% conditions varied slightly across the $16 and $32 endowments. These trials were assigned to their four canonical fairness levels and coded as 0.05, 0.10, 0.25, or 0.50.

We estimated participant-specific fairness slopes using binomial generalized linear mixed-effects models with a logit link. For participant i on trial j, the model was

$$\text{logit\{P(Yᵢⱼ = 1)\} = (β₀ + b₀ᵢ) + (β₁ + b₁ᵢ)Fᵢⱼ}$$

where Yᵢⱼ denotes the accept/reject decision and Fᵢⱼ denotes the canonical fairness level. The fixed coefficient β₁ estimated the sample-level association between fairness and acceptance. The random coefficient b₁ᵢ allowed this association to vary across participants, while b₀ᵢ allowed baseline acceptance to vary across participants. Models were estimated with the glmer function from the lme4 package in R using the formula

$$\text{accept \textasciitilde fairness + (1 + fairness | participant)}$$

and a binomial error distribution.

For each model, the participant-specific fairness slope was the sum of the fixed fairness coefficient and that participant’s random-slope deviation, β̂₁ + b̂₁ᵢ. These conditional slopes were expressed on the log-odds scale and were partially pooled toward the sample-level estimate. Separate models produced a social slope from human-partner trials, a nonsocial slope from computer-partner trials, and a combined slope from all trials. The combined slope was estimated directly from the all-trials model rather than calculated as the mean of the social and nonsocial slopes. Social-relative fairness sensitivity was calculated as

$$\text{Dᵢ = Sᵢ - Nᵢ}$$

where Sᵢ and Nᵢ denote participant i’s social and nonsocial slopes, respectively. Larger positive slopes indicated that acceptance increased more steeply as offers became fairer; larger positive difference scores indicated greater fairness sensitivity for human than computer partners.

### Neuroimaging Data Preprocessing

All imaging data were first converted to BIDS format using HeuDiConv (Halchenko et al., 2024) prior to preprocessing. The results reported in this manuscript are based on data processed with fMRIPrep 24.1.1 (Esteban et al., 2019; Markiewicz et al., 2026), which builds on the Nipype framework (version 1.8.6, Esteban et al., 2025; Gorgolewski et al., 2011). Portions of the preprocessing description below are adapted from the standard fMRIPrep boilerplate and edited for clarity and relevance to the present study.

#### Anatomical data preprocessing.

For each participant, the T1w image was corrected for intensity non-uniformity (INU) with N4BiasFieldCorrection (Tustison et al., 2010), distributed with ANTs 2.5.3 (Avants et al., 2008), and used as T1w-reference throughout the workflow. The T1w-reference was then skull-stripped with a Nipype implementation of the antsBrainExtraction.sh workflow (from ANTs), using OASIS30ANTs as target template. Brain tissue segmentation of cerebrospinal fluid (CSF), white-matter (WM) and gray-matter (GM) was performed on the brain-extracted T1w using FAST (FSL 6.0.4, Zhang et al., 2001). Volume-based spatial normalization to two standard spaces (MNI152NLin6Asym, MNI152NLin2009cAsym) was performed through nonlinear registration with antsRegistration (ANTs 2.5.3), using brain-extracted versions of both T1w reference and the T1w template. The following templates were selected for spatial normalization and accessed with TemplateFlow (24.2.0, Ciric et al., 2022): FSL’s MNI ICBM 152 non-linear 6th Generation Asymmetric Average Brain Stereotaxic Registration Model (TemplateFlow ID: MNI152NLin6Asym, Evans et al., 2012), ICBM 152 Nonlinear Asymmetrical template version 2009c (TemplateFlow ID: MNI152NLin2009cAsym, Fonov et al., 2009).

#### Functional data preprocessing.

For all functional runs per participant, the following preprocessing was performed. First, a reference volume was generated from the shortest echo of the BOLD run, using a custom methodology of fMRIPrep, for use in head motion correction. Head-motion parameters with respect to the BOLD reference (transformation matrices, and six corresponding rotation and translation parameters) are estimated before any spatiotemporal filtering using mcflirt (FSL, Jenkinson et al., 2002). The estimated fieldmap was then aligned with rigid-registration to the target EPI (echo-planar imaging) reference run. The field coefficients were mapped onto the reference EPI using the transform. The BOLD reference was then co-registered to the T1w followed by flirt (FSL, Jenkinson & Smith, 2001) with the boundary-based registration (Greve & Fischl, 2009) cost-function. Co-registration was configured with six degrees of freedom. Several confounding time-series were calculated based on the preprocessed BOLD: framewise displacement (FD), DVARS and three region-wise global signals. FD was computed using two formulations following Power (absolute sum of relative motions, Power et al., 2014) and Jenkinson (relative root mean square displacement between affines, Jenkinson et al., 2002). FD and DVARS are calculated for each functional run, both using their implementations in Nipype (following the definitions by Power et al., 2014). The three global signals are extracted within the CSF, the WM, and the whole-brain masks.

Additionally, a set of physiological regressors were extracted to allow for component-based noise correction (CompCor, Behzadi et al., 2007). Principal components are estimated after high-pass filtering the preprocessed BOLD time-series (using a discrete cosine filter with 128 s cut-off) for the two CompCor variants: temporal (tCompCor) and anatomical (aCompCor). tCompCor components are then calculated from the top 2% variable voxels within the brain mask. For aCompCor, three probabilistic masks (CSF, WM and combined CSF+WM) are generated in anatomical space. The implementation differs from that of Behzadi et al. in that instead of eroding the masks by 2 pixels on BOLD space, a mask of pixels that likely contain a volume fraction of GM is subtracted from the aCompCor masks. This mask is obtained by thresholding the corresponding partial volume map at 0.05, and it ensures components are not extracted from voxels containing a minimal fraction of GM. Finally, these masks are resampled into BOLD space and binarized by thresholding at 0.99 (as in the original implementation). Components are also calculated separately within the WM and CSF masks. For each CompCor decomposition, the k components with the largest singular values are retained, such that the retained components’ time series are sufficient to explain 50 percent of variance across the nuisance mask (CSF, WM, combined, or temporal). The remaining components are dropped from consideration. The head-motion estimates calculated in the correction step were also placed within the corresponding confounds file. All resamplings can be performed with a single interpolation step by composing all the pertinent transformations (i.e. head-motion transform matrices, susceptibility distortion correction when available, and co-registrations to anatomical and output spaces). Gridded (volumetric) resamplings were performed using nitransforms, configured with cubic B-spline interpolation. Many internal operations of fMRIPrep use Nilearn 0.10.4 (Abraham et al., 2014), mostly within the functional processing workflow. For more details of the pipeline, see the section corresponding to workflows in fMRIPrep's documentation (https://fmriprep.readthedocs.io/en/latest/workflows.html).

Further, we applied spatial smoothing with a 5 mm full-width at half-maximum (FWHM) Gaussian kernel to all functional data using FMRI Expert Analysis Tool (FEAT), version 6.0.2, part of FSL (FMRIB’s Software Library, www.fmrib.ox.ac.uk/fsl). Non-brain removal using BET (S. M. Smith, 2002) and grand mean intensity normalization of the entire 4D dataset by a single multiplicative factor were also applied.

### Echo-Time Dependent Denoising

TE-dependence analysis was performed on input data using the tedana workflow (DuPre et al., 2021). An initial mask was generated from the first echo using nilearn's compute_epi_mask function. An adaptive mask was then generated using the dropout method(s), in which each voxel's value reflects the number of echoes with 'good' data. A two-stage masking procedure was applied, in which a liberal mask (including voxels with good data in at least the first echo) was used for optimal combination, T2*/S0 estimation, and denoising, while a more conservative mask (restricted to voxels with good data in at least the first three echoes) was used for the component classification procedure. A monoexponential model was fit to the data at each voxel using nonlinear model fitting in order to estimate T2* and S0 maps, using T2*/S0 estimates from a log-linear fit as initial values. For each voxel, the value from the adaptive mask was used to determine which echoes would be used to estimate T2* and S0. In cases of model fit failure, T2*/S0 estimates from the log-linear fit were retained instead. Multi-echo data were then optimally combined using the T2* combination method (Posse et al., 1999). Principal component analysis based on the PCA component estimation with a Moving Average (stationary Gaussian) process (Li et al., 2007) was applied to the optimally combined data for dimensionality reduction. The following metrics were calculated: kappa, rho, countnoise, countsigFT2, countsigFS0, dice_FT2, dice_FS0, signal-noise_t, variance explained, normalized variance explained, d_table_score. Kappa (kappa) and Rho (rho) were calculated as measures of TE-dependence and TE-independence, respectively. A t-test was performed between the distributions of T2*-model F-statistics associated with clusters (i.e., signal) and non-cluster voxels (i.e., noise) to generate a t-statistic (metric signal-noise_z) and p-value (metric signal-noise_p) measuring relative association of the component to signal over noise. The number of significant voxels not from clusters was calculated for each component. Independent component analysis was then used to decompose the dimensionally reduced dataset. The following metrics were calculated: countnoise, countsigFS0, countsigFT2, d_table_score, dice_FS0, dice_FT2, kappa, normalized variance explained, rho, signal-noise_t, variance explained. Kappa (kappa) and Rho (rho) were calculated as measures of TE-dependence and TE-independence, respectively. A t-test was performed between the distributions of T2*-model F-statistics associated with clusters (i.e., signal) and non-cluster voxels (i.e., noise) to generate a t-statistic (metric signal-noise_z) and p-value (metric signal-noise_p) measuring relative association of the component to signal over noise. The number of significant voxels not from clusters was calculated for each component. Next, component selection was performed to identify BOLD (TE-dependent) and non-BOLD (TE-independent) components using a decision tree.

This workflow used numpy (Van Der Walt et al., 2011) scipy (Virtanen et al., 2020), pandas (McKinney, 2010; The pandas development team, 2024), scikit-learn (Pedregosa et al., 2011), nilearn (Nilearn contributors et al., 2026), bokeh (Bokeh Development Team, 2018), matplotlib (Hunter, 2007), and nibabel (Brett et al., 2019). This workflow also used the Dice similarity index (Dice, 1945; Sorenson, 1948).

### Creation of B0 Field Maps

We created static B0 field maps from the phase information in our multi-echo functional MRI (ME-fMRI) echo-planar imaging (EPI) data using the Multi-Echo DIstortion Correction (MEDIC) algorithm (version 0.1.1; https://github.com/vanandrew/warpkit) (Van et al., 2023). This method estimates voxelwise B0 inhomogeneity by modeling the phase of the MRI signal as a linear function of echo time (Jezzard & Balaban, 1995). To remove channel-specific phase offsets introduced during multi-coil image reconstruction, we applied Multi-Channel Phase Combination using 3D Simultaneous estimation (MCPC-3D-S), which derives a zero-time phase offset based on the unwrapped phase difference between the first two echoes (Eckstein et al., 2018). Phase unwrapping was then performed using Rapid Open-source Minimum-spanning-tree phase unwrapping (ROMEO), which jointly unwraps all echoes under the constraint that phase accumulates linearly with echo time, thereby improving stability and accuracy (Dymerska et al., 2021). The resulting unwrapped phase data were fitted using a magnitude-weighted least-squares approach to compute off-resonance field maps in units of hertz, which were then transformed to anatomical space for use in susceptibility distortion correction.

### First-Level Nuisance Regressors

First-level activation and connectivity models included nuisance regressors to account for motion and physiological noise. These included the six rigid-body motion parameters, the first six aCompCor components, non-steady-state volumes, and framewise displacement. TEDANA-derived nuisance components classified as non-BOLD were included as additional regressors to further account for residual non-BOLD variance. High-pass filtering was implemented with a 128 s cut-off using discrete cosine basis functions.
